# STAT3 promotes melanoma metastasis by CEBP-induced repression of the MITF pigmentation pathway

**DOI:** 10.1101/422832

**Authors:** Alexander Swoboda, Robert Soukup, Katharina Kinslechner, Bettina Wingelhofer, David Schörghofer, Christina Sternberg, Ha T. T. Pham, Maria Vallianou, Jaqueline Horvath, Dagmar Stoiber, Lukas Kenner, Lionel Larue, Valeria Poli, Friedrich Beermann, Takashi Yokota, Stefan Kubicek, Thomas Krausgruber, André F. Rendeiro, Christoph Bock, Rainer Zenz, Boris Kovacic, Fritz Aberger, Markus Hengstschläger, Peter Petzelbauer, Mario Mikula, Richard Moriggl

## Abstract

Metastatic melanoma is hallmarked by its ability to switch oncogenic MITF expression. Here we tested the impact of STAT3 on melanoma onset and progression in association with MITF expression levels. We established a mouse melanoma model for deleting *Stat3* specifically in melanocytes with specific expression of human hyperactive *NRAS*^*Q61K*^ in an *Ink4a* deficient background. Mice with tissue specific *Stat3* deletion showed an early onset of disease, but displayed significantly diminished lung metastases. Whole genome expression profiling also revealed a reduced invasion phenotype, which was functionally confirmed in 3D melanoma model systems. Notably, loss or knockdown of STAT3 in mouse or human cells resulted in up-regulation of MITF and induction of cell proliferation. Mechanistically we show that STAT3 induced CEBPa/b expression was sufficient to suppress MITF transcription. Epigenetic analysis by ATAC-seq confirmed that STAT3 enabled CEBPa/b binding to the *MITF* enhancer region thereby silencing it. We conclude that STAT3 is a metastasis driver in melanoma able to antagonize the *MITF* oncogene via direct induction of CEBP family member transcription facilitating RAS-RAF-driven melanoma metastasis.

**List of Abbreviations**

ATAC-seq, Assay for Transposase-Accessible Chromatin using sequencing; CEBP, CAAT Box Enhancer Binding Protein; CRE, Cre recombinase; EGF, Epidermal Growth Factor; GEO, Gene Expression Omnibus; GSEA, Gene Set Enrichment Analysis; HSC70, Heat Shock 70 kDa protein; IHC, Immunohistochemistry; IL-6, Interleukin-6; JAK, Janus Kinase; MITF, Microphthalmia-Associated Transcription Factor; NSG, NOD.Cg-Prkdc^scid^ Il2rg^tm1Wjl^/SzJ mice; OSM, Oncostatin M; PDGF, Platelet-Derived Growth Factor; pS, phosphoserine; pY, phosphotyrosine; RAS, Rat Sarcoma; RAF, Rapidly Accelerated Fibrosarcoma; RTK, Receptor Tyrosine Kinase; RT-PCR, Reverse Transcription Polymerase Chain Reaction; S100b, Calcium Binding Protein S100 beta; shRNA, short hairpin RNA; SOX10, Sex Determining Region Y-10; STAT, Signal Transducer and Activator of Transcription; TCGA, The Cancer Genome Atlas; TMA, Tissue Micro Array

## Introduction

Melanoma is a very aggressive form of skin cancer, with >76.000 new cases diagnosed annually in the US ^1^. Stage I and stage II lesions can be successfully removed by surgery, but metastasized melanomas are challenging to treat, leading to an estimated 10.000 deaths in the US annually ^1^. The current prognosis of advanced melanoma remains poor despite the success of immune- and targeted therapy ^2^. This phenomenon can be attributed to the plasticity of melanoma, which describes the ability of melanoma cells to switch multiple times from a proliferative to an invasive state without the need for additional mutations ^3,4^. A key player involved in phenotype switching of melanoma is the Microphthalmia-associated Transcription Factor (MITF). MITF is essential for melanocyte development, homeostasis and pigmentation response ^5,6^. This oncoprotein controls differentiation, survival and proliferation. High MITF expression marks melanoma cells with a proliferative, but non-invasive phenotype. In contrast, melanoma cells expressing low MITF protein represent increased invasive and metastatic capacity ^3,7-9^. Overall, the regulation of MITF by *NRAS* and *BRAF* oncogenes in context of STAT3 signaling is unclear ^7,10^.

Interestingly, about 50% of melanoma patients display prominent and enhanced tyrosine phosphorylation of Signal Transducer and Activator of Transcription 3 (STAT3) catalyzed via JAK, SRC or growth factor tyrosine kinase family members ^11-14^. So far, reports on inhibition of STAT3 by siRNA or expression of a dominant negative form of STAT3 in xenograft models have postulated an oncogenic role of STAT3 in melanoma progression, but detailed genetic studies are still missing ^12,15^. Contrary, treatment with Oncostatin M (OSM) or IL-6 resulted in phosphotyrosine-mediated STAT3 (pYSTAT3) activation and decreased proliferation in melanoma cell line studies, suggesting a tumor suppressor role for STAT3 in melanoma progression ^14,16^. In prostate, lung and colorectal carcinomas, tumor suppressive roles were associated with STAT3 function depending on the mutational context or STAT1 expression levels, a known heterodimerization partner/balancer of oncogenic STAT3 ^17-20^. A specific role for STAT3 in melanoma has not yet been established and it is not clear whether STAT3 plays a role in phenotype-switching towards invasive melanoma.

Here, we utilized a genetic model of spontaneous melanoma formation. To closely mimic human melanoma, transgenic mice have been used carrying melanocyte-specific expression of the *NRAS*^*Q61K*^ oncogene in an INK4A-deficient background ^21^. Additionally, the mouse model allows for conditional melanocyte-specific deletion of *Stat3* ^22^. We show that melanoma lacking *Stat3* expression have an accelerated tumor onset in vivo and exert higher proliferation rate. In contrast, metastasis formation from *Stat3* knockout primary tumors was severely impaired. STAT3 function was evaluated mechanistically using tumor derived cell lines, where we performed whole genome expression analysis combined with ATAC-seq profiling. We found, that STAT3 antagonizes MITF expression through elevated expression of CAAT Box Enhancer Binding Protein (CEBP) family members. Remarkably, data mining of melanoma patient samples also revealed a negative correlation of CEBPa/b with MITF. STAT3 knockdowns in human melanoma confirmed these observations. We conclude that STAT3 is associated with melanoma metastasis and MITF is associated with melanoma growth. Therefore, concurrent low STAT3 and high MITF expression marks worsened prognosis for melanoma patients.

## Results

### Loss of STAT3 enhanced pigmentation, enabled earlier tumor onset, but reduced metastasis

To study the effect of STAT3 loss in melanoma we used a mouse model that allows the conditional deletion of the *Stat3* gene by C*re-loxP* technology. We investigated how deletion of the *Stat3* gene influences progression of melanoma in a genetic mouse model driven by melanocyte specific dominant-active human *NRAS*^*Q61K*^. Furthermore, the *INK4a* tumor suppressor locus was deleted, hence the model closely mimics human cutaneous melanoma progression. Melanoma that carry *NRAS*^*Q61K*^, lost p16^INK4A^ and p19^ARF^ and deleted *Stat3* (Tyr∷NRAS^Q61K^Ink4a^-/-^Stat3^flox/flox^Tyr∷Cre) are further termed Stat3^Δ^, control melanoma expressing STAT3 are termed Stat3^fl^ (Fig. 1 A). Melanocyte specific CRE expression was described to recombine *loxP* sites from E10.5 onwards in development ^23^. Hence, STAT3 was lost in mouse skin melanomas as evaluated by immunohistochemistry (IHC, Fig. 1 B). PCNA staining revealed that primary Stat3^Δ^ tumors displayed increased cell cycle activity compared to Stat3^fl^ tumors (Fig. 1 C). In line with this, we observed significantly accelerated tumor onset in the Stat3^Δ^ group (Fig. 1 D). Notably, despite earlier onset of disease, the overall survival was comparable (Fig. S1 A). Stat3^Δ^ tumors appeared more pigmented compared to Stat3^fl^ tumors and Melanin measurements of isolated and cultured melanocytes confirmed this finding (Fig. 1 E). Since loss of *Stat3* could impact expression of the nearby *Stat5* locus, we tested STAT5 expression and localization by antibody staining but no compensation occurred (Fig. S1 B and C).

**Figure 1.**
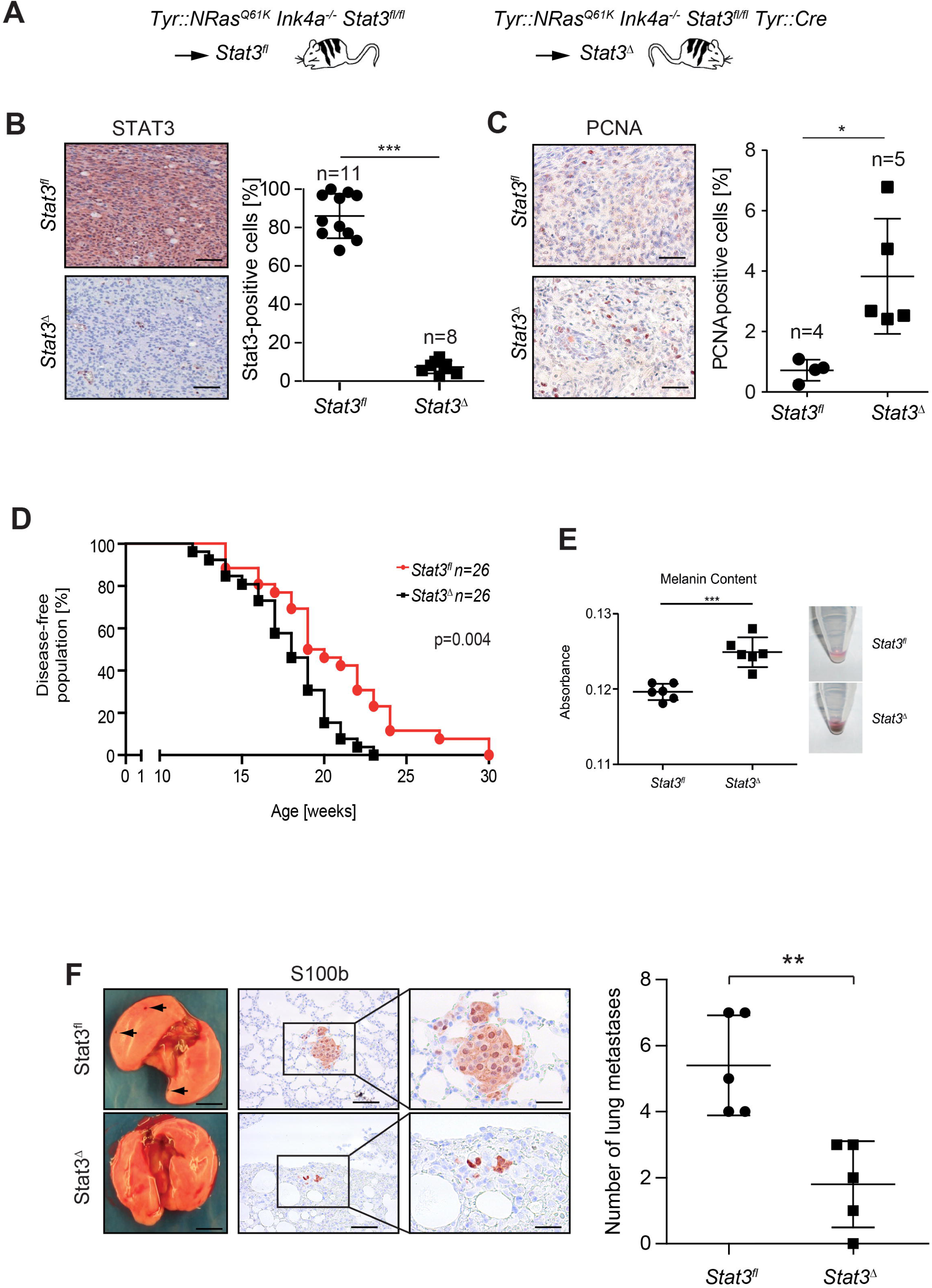
STAT3 knockout in melanoma induced earlier tumor onset, but reduced metastasis. (A) Mice containing a constitutively active *NRAS* gene controlled by the *tyrosinase* promoter and a deletion of the *Ink4a* locus were crossed to mice harboring a *floxed Stat3* locus termed “Stat3^fl^”. Mice additionally expressing *Cre* recombinase by a *tyrosinase* promoter were termed “Stat3^Δ^". (B+C) IHC evaluation of total STAT3 and PCNA in primary melanoma of Stat3^fl^ and Stat3^Δ^ mice. Scale bars, 50 μm. (D) Kaplan–Meier plot, showing prolonged disease-free survival, defined as time before palpable tumors occur, of Stat3^fl^ mice compared to the Stat3^Δ^ group. (E) Tumor derived mouse melanoma cell lines were seeded in 6-wells and cultured under standard conditions. After 48 hours of culturing, melanin content of the supernatant was measured by absorption at 410 nm. Results represent six independent measurements. Data were analyzed by Student’s t-test. (F) Representative lungs, arrows indicate melanoma metastases (left); S100b staining (middle) and metastasis quantification (right) of age matched lung samples of Stat3^fl^ and Stat3^Δ^ mice. Scale bar (left), 3 mm, scale bar (middle), 60 µm, scale bar (right), 150□μm. Data in B, C and F were analyzed by unpaired 2-tailed Student’s t-test \*\*\**P*<0.001, **P*<*0.01, \**P*<0.1. Logrank (Mantel-Cox) tests were used for the analysis of the Kaplan–Meier plot in D.

Next, lungs of 40 week old mice were stained for S100b to visualize micrometastases. Importantly, *Stat3* deletion significantly reduced the overall number of metastatic lung colonies (Fig. 1 F). These findings indicate that albeit tumor growth was accelerated in the Stat3^Δ^ group, *Stat3* expression enhanced the metastatic process. Next, we investigated the mechanism responsible for reduced metastasis of Stat3^Δ^ melanomas.

### Expression profiling revealed MITF pathway induction upon loss of STAT3

We isolated tumor cells from melanoma positive lymph nodes of the Stat3^fl^ and Stat3^Δ^ mice and selected for melanoma cells by continuous culturing for 10 passages. Cells were uniformly positive for the melanoma marker S100b (Fig. S1 D). They showed basal or enhanced STAT3 activity after IL-6 stimulation, while Stat3^Δ^ cells showed complete loss of STAT3 expression as measured by STAT3 tyrosine and serine phosphorylation using specific antibody detection by Western blotting (Fig. 2 A). Then we performed Affymetrix whole transcriptome microarray mRNA analysis followed by gene set enrichment analysis (GSEA) under basal growth conditions or stimulation with murine IL-6 or OSM. Loss of *Stat3* resulted in significant reduction of *Stat3* target gene expression (Fig. 2 B). Importantly, Stat3^Δ^ cells displayed augmented MITF pathway activation, which was amongst the top up-regulated pathways (Fig. 2 B). To our surprise, cytokine stimulation neither influenced the overall MITF pathway regulation by activated STAT3 (Fig. 2 C, D and Fig. S2), nor changed the overall gene expression after two hours of stimulation significantly. We conclude that low pYSTAT3 is actively engaged for broad *Stat3* target gene transcription by oncogenic RAS-RAF signaling to reprogram melanoma cells, independent of a cytokine stimulus.

**Figure 2.**
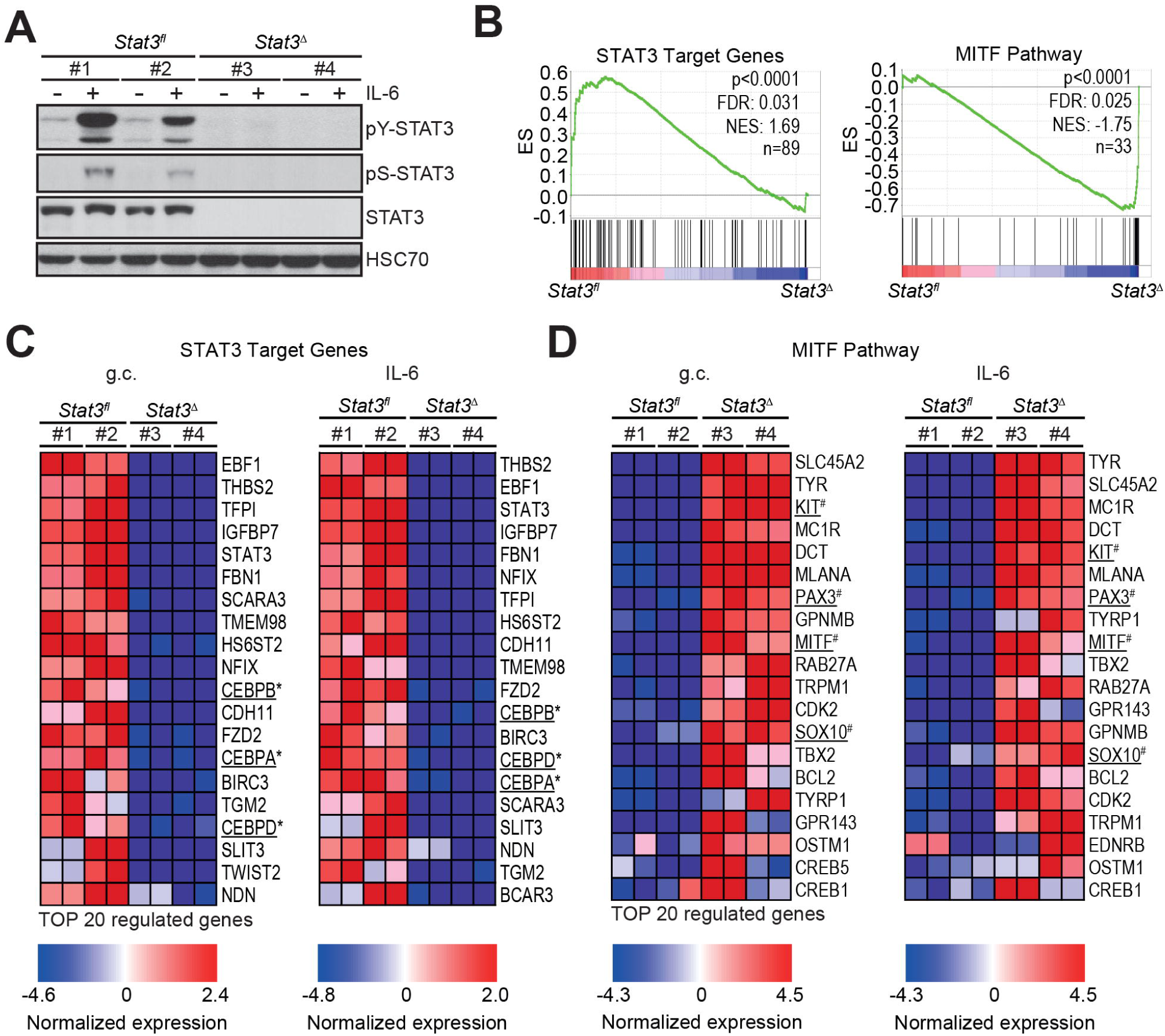
Loss of *Stat3* induced MITF pathway in melanoma cells. (A) Representative Western blot showing total STAT3 levels and IL-6 stimulated (20 ng/ml, 30 minutes) STAT3 phosphorylation at Y705 or S727 in two representative cell lines derived from either Stat3^fl^ or Stat3^Δ^ tumors. HSC70 served as loading control. (B) STAT3 targets of 89 genes (Table S4) and a MITF pathway set comprised of 33 MITF targets (Table S5) were evaluated by gene-set enrichment analysis (GSEA) during normal growth in cell culture. (C) Heatmaps of the top 20 down-regulated genes of the STAT3 gene-set shown in B in Stat3^Δ^ cells during normal growth conditions (g.c.) (left) and after 2 hours of stimulation with IL-6 (right). (D) Heatmaps of the top 20 up-regulated genes of the MITF gene-set shown in B in Stat3^Δ^ cells during normal g.c. (left) and after 2 hours of stimulation with IL-6 (right).

### Invasion and EMT-like phenotype are suppressed upon loss of *Stat3*

Gene set enrichment analysis, including proliferative and invasive melanoma signatures, were performed to identify STAT3 related phenotypes (Fig. 3 A). We found that Stat3^Δ^ cells resembled MITF-driven proliferative gene signatures, which were identified in two independent large melanoma cohort studies ^3,9^. In contrast, Stat3^fl^ cells are closely related to the described invasive signatures. Performing *in vitro* assays, we found enhanced 3D-proliferation, but abrogated invasion and migration in Stat3^Δ^ cells (Fig. 3 B and C). Our data implies that STAT3 fulfills an important function in RAS-RAF-transformed melanoma promoting invasion and migration. Tumor xenografts of Stat3^Δ^ cells were more compact, with higher cellular density and showed little to no invasion of epidermal tissue (Fig. 3 D). In contrast Stat3^fl^-derived tumors were de-differentiated with fibroblastoid morphology and showed high invasion into murine epidermis. Tumor formation and growth rates of the two melanoma groups after xenotransplantation were comparable (Fig. S3 A).

**Figure 3.**
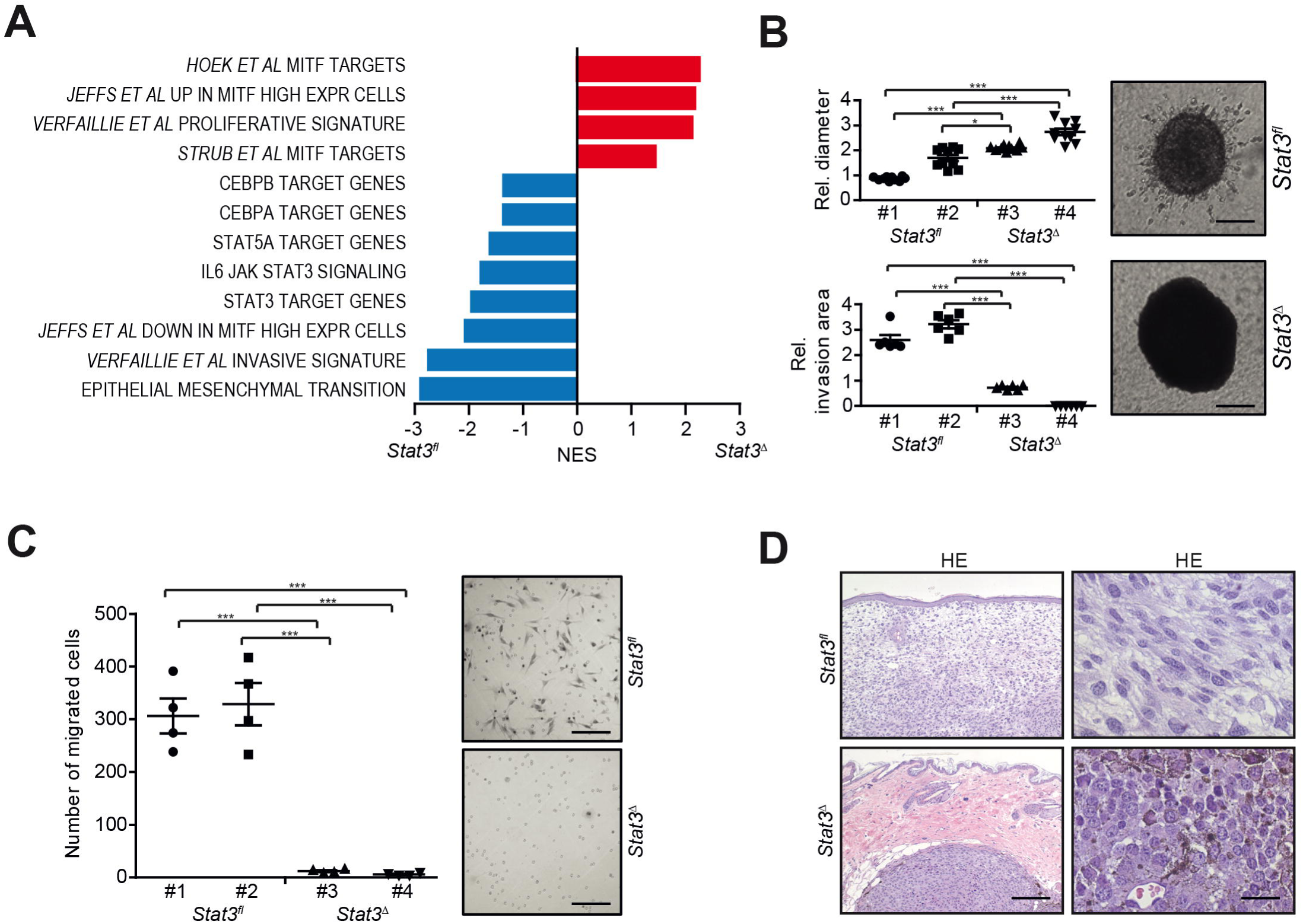
Transcriptome analysis and functional testing revealed abrogated invasion and increased proliferation after STAT3 knockout. (A) Normalized enrichment scores (NES) calculated for target gene signatures related to STAT3 and MITF derived from the GSEA database or melanoma gene signatures derived from publications. Gene-sets indicate a more proliferative and less invasive phenotype. Cutoff values: False discovery rate: <5%, *p*<0.05. (B) 3D-proliferation and sphere invasion assay of Stat3^fl^ and Stat3^Δ^ melanoma cells into a collagen gel (n=10/group). Stat3^Δ^ cells proliferate faster (upper panel) and have abrogated invasive capabilities (n=6/group) (lower panel). Two representative spheres are displayed. Scale bar, 200 μm. (C) Transwell migration assay of Stat3^fl^ and Stat3^Δ^ melanoma cells (n=4). STAT3 deletion leads to abrogated migration. Scale bar, 100 μm. (D) Representative HE stainings of tumors formed from Stat3^fl^ and Stat3^Δ^ cells xenografted into NSG mice. Subcutaneously injected tumors of the Stat3^fl^ group invaded the dermis, while tumors of the Stat3^Δ^ group have a compact structure with even borders that did not invade into the dermis. Scale bar (left), 300□μm, scale bar (right), 20□μm.

Beside the prominent MITF pathway expression, we found a set of five important de-regulated receptor tyrosine kinases: Three of them, platelet-derived growth factor receptor alpha and beta (PDGFRa/b) and epidermal growth factor receptor (EGFR). These are known targets of STAT3 and accordingly displayed decreased mRNA expression in the Stat3^Δ^ group (Fig. 4 A and Figure S3 B). For the two other receptor tyrosine kinases, *Met* and *cKit*, expression was in contrast increased. Both are oncogenes and act as important inducers of MITF pathway activity and Melanin production in normal melanocytes. Accordingly, PDGFRb and EGFR were highly expressed in immunostainings of Stat3^fl^ primary mouse melanomas, whereas Stat3^Δ^ tumors showed increased expression of MET and cKIT (Fig. 4 B and C).

**Figure 4.**
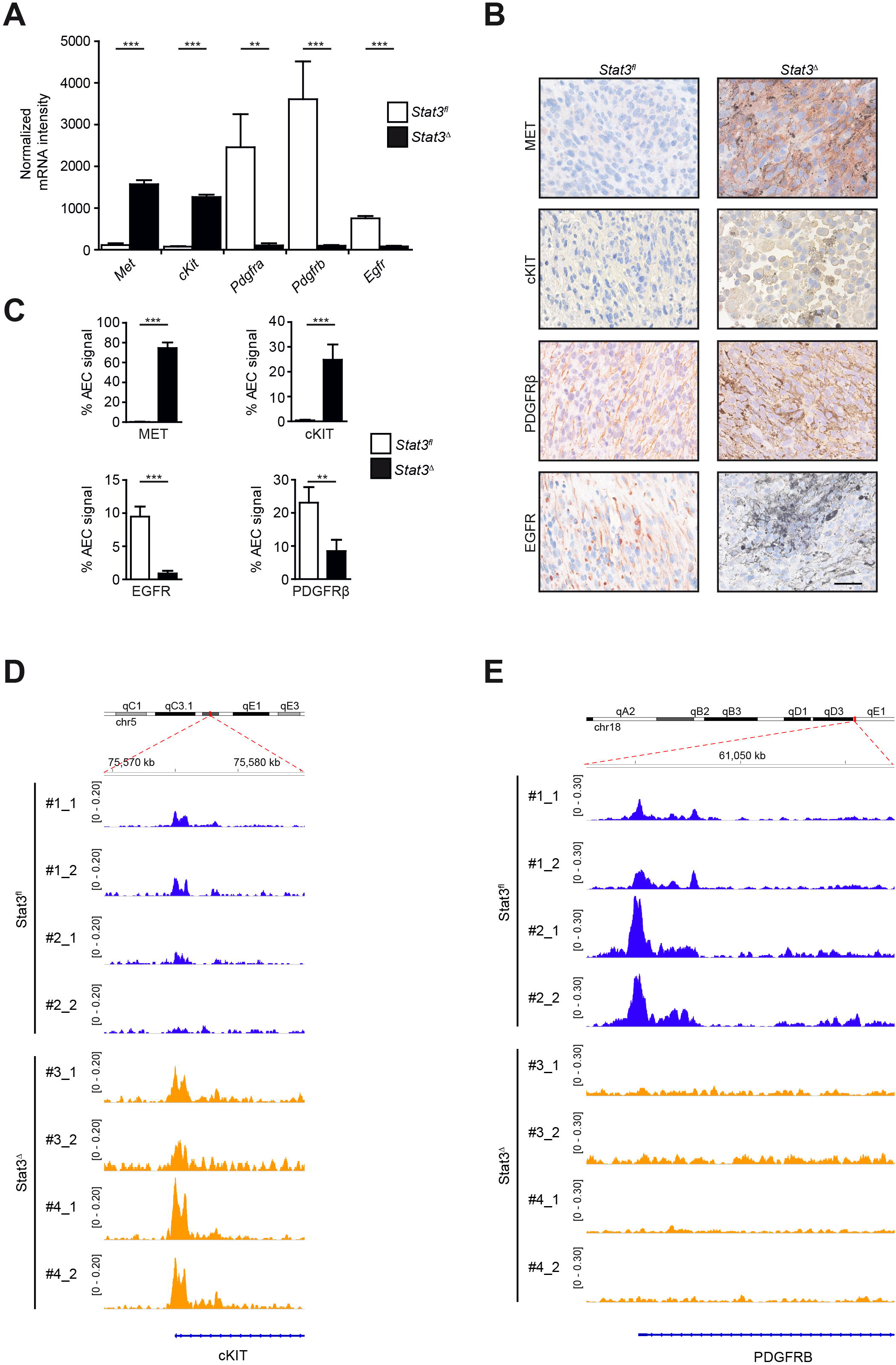
Expression of receptor tyrosine kinases display a YIN/YANG dualism corresponding to STAT3/MITF interplay. (A) The total mRNA levels of a set of significantly regulated RTK related to STAT3 and MITF signaling after normalization of the whole transcriptome expression screen comparing the expression levels between the Stat3^fl^ and the Stat3^Δ^ group. (B) Representative images of IHCs of RTK expression in tumors of xenografted Stat3^fl^ and Stat3^Δ^ mouse melanoma cell lines in NSG mice (C) and quantifications thereof (n=4, each). Red color depicts specific immunostaining and black/brown is related to pigmentation. Scale bar, 25□μm. (D+E) ATAC-seq signal intensities at the *cKit* and at the P*dgfrb* locus. Depicted in blue are STAT3^fl^ cell lines and in yellow STAT3^Δ^ cell lines all in technical duplicates.

To dissect the regulatory basis of the changes in the cell transcriptomes, we performed chromatin profiling using ATAC-seq with a focus on the *cKit* and *Pdgfrb* locus in Stat3^fl^ and Stat3^Δ^ melanoma cells. Chromatin accessibility at the *cKit* promoter region was increased only in Stat3^Δ^ and conversely the *Pdgfrb* promoter was only accessible in Stat3^fl^ cells (Fig. 4 D and E). This indicates, that the loss of STAT3 leads to epigenetic changes accompanied by the observed strong changes in gene expression patterns.

### Reciprocal expression between CEBPs and MITF

To investigate the mechanisms underlying the regulation of MITF by STAT3, we reasoned that STAT3 or its downstream targets could repress MITF. To test this hypothesis, we investigated regulatory elements of the proximal *Mitf* promoter region by ATAC seq. We found that Stat3^fl^ cells displayed chromatin accessibility in a regulatory element in close proximity of the *Mitf* gene which contained a CEBPa/b binding element (Fig. 5 A and B). Access at this site was specifically lost in Stat3^Δ^ cells. Contrary, Stat3^Δ^ cells displayed an increased accessibility at a regulatory element containing a SOX10 binding element that was absent in Stat3^fl^ cells arguing for a SOX10-mediated establishment of a stable *Mitf* ^high^ status (Fig. 5 B).

**Figure 5.**
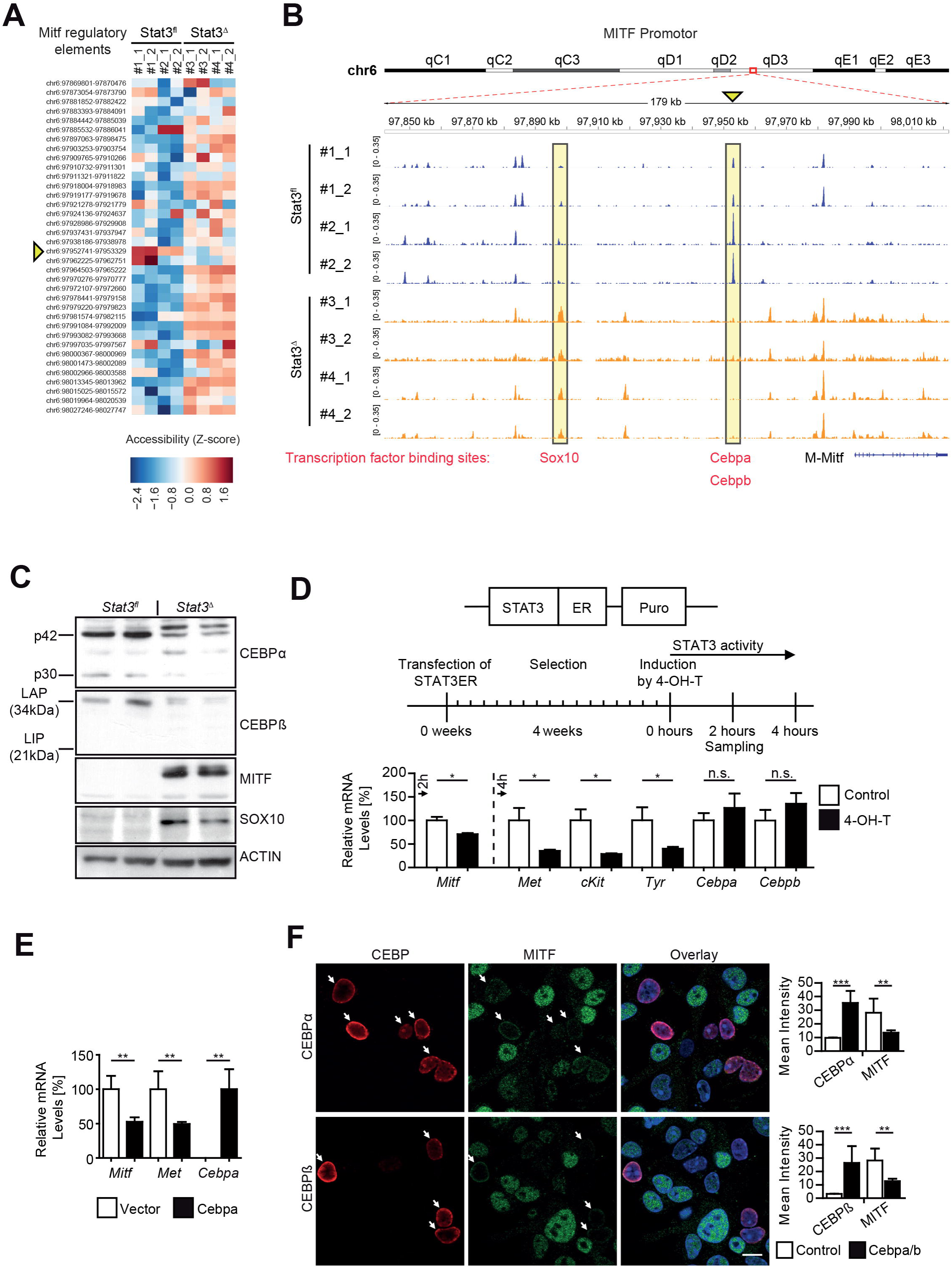
MITF expression depends on the STAT3 target *Cebpa* and *Cebpb*. (A) Heatmap displays the grade of accessible chromatin from MITF regulatory elements. Blue correlates with closed or less accessible chromatin and red with open or more accessible chromatin for binding factors. (B) ATAC-seq signal intensities at the M-MITF locus and the possible binding sites of SOX10 and CEBPa/b. Data mapped according to ChIP-Atlas. Depicted in blue are Stat3^fl^ cell lines and in yellow Stat3^Δ^ cell lines all in technical duplicates. (C) Western blots show increased expression of both the 42 kDa (p42) and the 30 kDa (p30) isoform of CEBPa/b and decreased expression of MITF and SOX10 in Stat3^fl^ cells in comparison to the Stat3^Δ^ group. (D) Lipofectamine transfection and stable selection via puromycin of Stat3^fl^ murine melanoma cells with a STAT3ER^T2^ construct that can be activated by 4-Hydroxytamoxifen (4-OH-T). MITF pathway is down-regulated after two to four hours after activation with 1 µM 4-OH-T. (E) *Mitf* and *Met* regulation after 24 hours transient *Cebpa* transfection of murine Stat3^Δ^ cells by lipofectamine. (F) Co-immunofluorescence for CEBPa or CEBPb and MITF expression after transient *Cebpa* or *Cebpb* transfection by lipofectamine for 24 hours in Stat3^Δ^ murine melanoma cells. Scale bar, 5□µm. Quantification of cells expressing CEBPa or CEBPb (n=4-10). Data in B, C and D were analyzed by unpaired 2-tailed Student’s t-test \**P*<0.05, \*\**P*<0.01, \*\*\**P*<0.001.

The CEBP transcription factors are well known downstream targets of STAT3 and we could identify strong inhibition of *Cebpa/b/d* mRNA expression after *Stat3* knockout (Fig. 2 C, underlined). Also on protein level we found CEBPa/b proteins to be significantly expressed in a STAT3-dependent way and we could confirm reciprocal expression with MITF and SOX10 (Fig. 5 C). We introduced a 4-OH-tamoxifen inducible STAT3ER^T2^ construct ^24^ into Stat3^Δ^ murine melanoma cells. Addition of 4-OH-tamoxifen led to an increased expression of *Cebpa*/*b* mRNA as measured by RT-PCR analysis (Fig. 5 D). Expression of the STAT3ER^T2^ fusion protein was confirmed by Western blotting (Fig. S3 C). Induced STAT3 activity led to slightly increased CEBP expression and a reciprocal down-regulation of MITF pathway associated genes (*Mitf, Met, cKit, Tyr* were all significantly repressed upon 4-OH-tamoxifen induced STAT3 activation) (Fig. 5 D). Importantly, exogenous *Cebpa* expression in Stat3^Δ^ melanoma cells and subsequent mRNA quantification lead to a significant reduction in *Mitf* and *Met* expression, altering levels of these genes to approximately 50% of control cells (Fig. 5 E). In line with these results, murine and human melanoma cells devoid of STAT3, but ectopically overexpressing CEBPa or CEBPb, showed reduced MITF protein expression which validated the above ATAC seq findings (Fig. 5 F and Figure S4 and S5). We conclude that STAT3-regulated CEBP proteins are responsible for repression of MITF at mRNA and protein levels.

To test, whether our findings also apply for human melanoma cell lines, we performed stable lentiviral shRNA mediated knockdown of *Stat3* in 451Lu, WM793B and WM35 cells. Evaluation of the knockdown was performed via RT-PCR and Western blot (Fig. 6 A and C). Consistent with our murine data, we observed up-regulation of *MITF, MET, SOX10* and *TYR* on mRNA level, as well as MITF and SOX10 up-regulation on protein level, upon *Stat3* knockdown (Fig. 6 B and C). Furthermore, cells devoid of *Stat3* showed an increased proliferation as quantified by Ki67 positive cells (Fig. 6 D). To further validate and to address the clinical relevance of our findings, we evaluated publically available expression data sets from the Gene Expression Omnibus (GEO) and The Cancer Genome Atlas (TCGA) databases. This analysis revealed a strong negative correlation between *MITF* and *CEBPA* as well as between *MITF* and *CEBPB* expression (Fig. 6 E).

**Figure 6.**
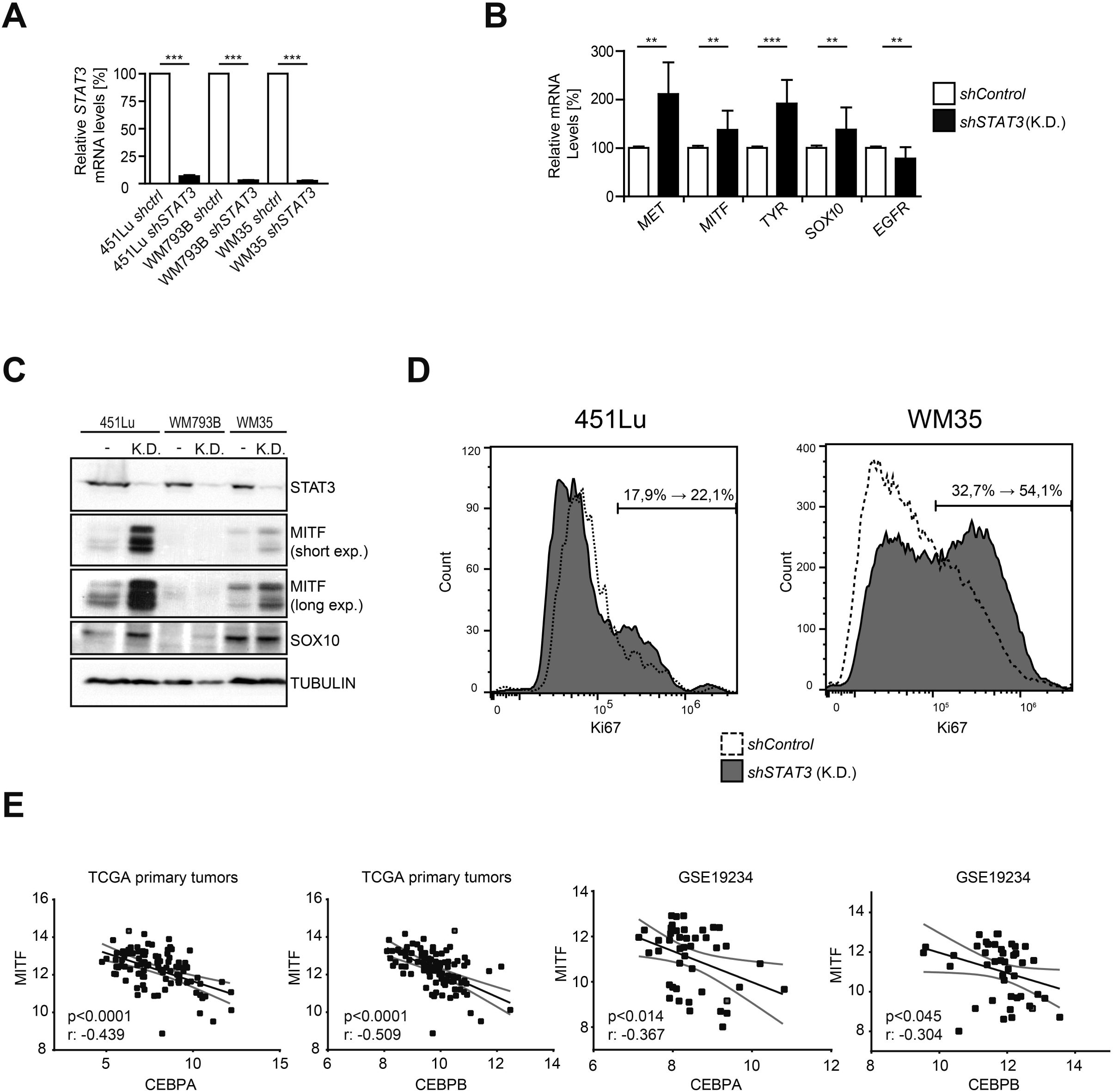
Human melanoma cells induce MITF and proliferation upon loss of STAT3. Three human melanoma cell lines were transduced with *STAT3* shRNA or a scrambled control by lentivirus and selected via puromycin resistance. (A) Evaluation of the *shSTAT3* RNA functionality by RT-PCR. (B) mRNA expression of MITF pathway members was up-regulated upon *STAT3* silencing in WM35 cells. (C) Evaluation of the *shSTAT3* RNA knockdown by Western blot for STAT3, SOX10 and MITF levels. Tubulin served as loading control. (D) FACS analysis of human melanoma cell lines stably expressing *shSTAT3* RNA or a scrambled control and stained against Ki67. (E) Data from the publically available “the cancer genome atlas – skin cutaneous melanoma (TCGA-SKCM)” and “GSE19234” human malignant melanoma patient datasets (Bogunovic, TCGA) were tested for correlations between *CEBPA* vs. *MITF* and *CEBPB* vs. *MITF*. All data concerning the Pearson correlation coefficients were calculated in SPSS (v21). Data in A and B were analyzed by unpaired 2-tailed Student’s t-test \**P*<0.05, \*\**P*<0.01, \*\*\**P*<0.001.

### Validation of STAT3 expression signatures in human clinical samples

To assess the clinical relevance of the relationship between MITF and CEBPa/b, we performed Kaplan-Meier survival analysis on publicly available datasets (GEO accession GSE19234 and TCGA data from Cutaneous Melanoma). Samples were sorted according to the expression level of *STAT3, CEBPA, CEBPB* and *MITF* to generate equal sized low- and high-risk groups (Fig. 7 A). The high-risk group was defined by high levels of *MITF* in combination with low *STAT3, CEBPA* and *CEBPB* levels (Fig. 7 B). To validate our findings in human patient samples we analyzed a cohort of 25 primary melanoma tumors and 96-130 melanoma metastases for STAT3 and MITF antibody mediated staining intensities. Primary tumors compared to metastases showed higher levels of STAT3 protein, while MITF levels were higher in metastases than in primary tumors (Fig. 7 C and Fig. S6). A representative example of consecutive sections of a primary melanoma and a metastatic lesion, stained for STAT3 and MITF is shown (Fig. 7 D).

**Figure 7.**
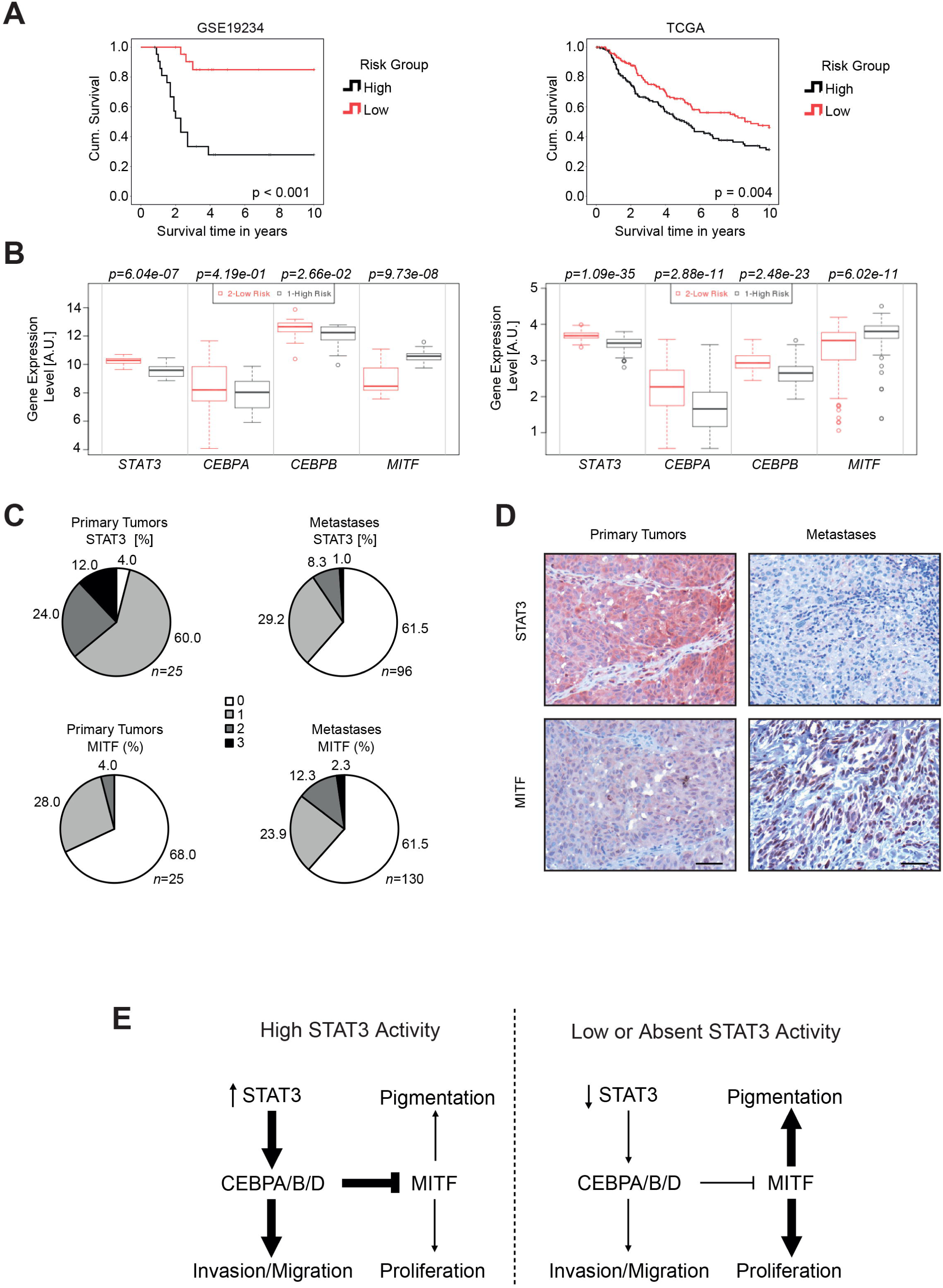
Human patients with STAT3^low^, CEBPA^low^, CEBPB^low^ and MITF^high^ signature show worsened clinical outcome. (A) Kaplan-Meier analysis was performed with the SurvExpress Tool and edited with SPSS v21 using datasets provided by *Bogunovic* et al. (44 samples, GSE19234) and the TCGA-SKCM database (335 samples). (B) For the analysis, samples were split at the median according to the prognostic index resulting in low- and high-risk groups. (C) Tissue microarray cohorts of 25 primary melanoma tumors and 96-130 melanoma metastases were stained for STAT3 and MITF levels. Decreasing STAT3 levels and increasing MITF levels were detected in metastases cohorts in comparison to the primary cohorts. For all graphs: 0=0-5%, 1=5-33%, 2=33-66%, 3=66-100% positive cells. (D) Consecutive sections of representative patient samples of a primary tumor and a metastasis stained against MITF and total STAT3. Scale bars, 50 μm. (E) Scheme depicting changes in transcription factor interaction governing the melanoma phenotype. In Stat3^fl^ mice, STAT3 pathway is active, driven by the mutated NRAS. STAT3 induces EGFR and PDGFRa/b creating a possible positive feedback on STAT3 signaling and establishing an EMT-like melanoma phenotype. When STAT3 activity is low or absent, mutant NRAS signaling is still active, but MITF is released from suppression by CEBP family members. Subsequently MITF target genes like *cKIT* and *MET* are up-regulated and these can also activate STAT3, promoting proliferation and survival. Additionally, MITF drives pigmentation by promoting a higher Melanin production. STAT3-CEBP expression stratifies patients into high and MITF expression into low risk groups, which could be of prognostic value.

In summary, we conclude that expression of CEBP family members depends on availability of STAT3. Upon *Stat3* deletion, *Cebpa, Cebpb* and *Cebpd* levels decreased, resulting in increased expression of *Mitf*, which abrogated invasion/migration and led to diminished metastasis formation (Fig. 7 E). In summary, the prognostic relevance is underlined by our findings that low STAT3 and high MITF expression exert a worsened prognosis for melanoma patients.

## Discussion

We identified STAT3 as a critical regulator to promote melanoma progression in a NRAS-driven mouse cutaneous melanoma model, devoid of INK4A tumor suppressors. STAT3 expression interferes with malignant melanoma growth through antagonizing MITF action via CEBPa/b/d family member induction. Analysis of publically available human mRNA expression datasets showed that high *MITF*, but low *STAT3* and *CEBPa/b* levels are associated with a worsened prognosis for patients with metastasis. Genetic loss or knockdown of STAT3 in mouse or human cells resulted in up-regulation of MITF and induction of cell proliferation. Tumor invasion and migration activity was abrogated upon STAT3 knockdown or loss, reminiscent to an EMT-like phenotype. Loss of STAT3 caused an earlier disease onset in mice consistent with a pronounced proliferative signature marked by augmented MITF expression. Rescuing Stat3^Δ^ melanoma cells by reactivation of the STAT3 pathway or by forced expression of CEBPa/b repressed *Mitf* mRNA production. We conclude that STAT3 is a major transcriptomic regulator able to establish persistent epigenetic changes and functionally promoting invasion and metastasis in melanoma.

Hence, targeting of STAT3 in melanoma therapy would drive tumors towards the melanocytic lineage and it would enhance proliferation. Targeting STAT3 only, could result in a worsened outcome, but targeting both oncogenic MITF and STAT3 could be beneficial for patients since both transcription factors control melanoma biology. Recent data showed that MITF inversely regulated the amount of EGFR, which was important to establish Vemuravenib resistance ^25^. Our data confirm this inverse association since we show that STAT3 induced MITF repression lead to EGFR and PDGFR accumulation. These findings display that MITF and STAT3 are tightly interwoven, best exemplified by tyrosine kinase receptor expression changes. While PDGFRa/b and EGFR were down-regulated in the Stat3^Δ^ group, MET and cKIT receptor showed strong up-regulation. These five receptor tyrosine kinases can all prominently activate STAT3 and their mode of action will depend on ligand-induced ligation via stromal components suggesting cell-cell interaction complexity, but also pointing towards the interdependence of the YIN/YANG STAT3/MITF dualism. MET and cKIT receptors are described as target genes of MITF and they could facilitate melanoma growth, invasion and angiogenesis ^26,27^. In addition, MET expression was associated with dependency of melanoma cells with a malignant phenotype ^28^. High cKIT levels are frequently found in benign skin lesions and primary melanomas associated with a less invasive phenotype ^29^. These reports support our finding that high MITF activity results in faster proliferation of melanomas to suppress migration and invasion. Interestingly, PDGFR expression in epithelial cells can be regarded as a marker for the EMT phenotype ^30,31^. The low response rate of patients towards anti-melanoma therapy was attributed to the persistence of a pronounced EMT-like phenotype ^32,33^. This could be due to the fact, that EMT in tumor patients is considered as a major factor in the development of resistance towards tyrosine kinase blockers ^34^ and for escaping PD-1 immunotherapy treatment ^35^. Our conclusion regarding STAT3 as a metastasis driver is paralleled by recent reports which describe STAT3 activity in melanoma cells as a driver for up-regulation of invasion-related genes and as a major factor in the transition towards a mesenchymal phenotype during EMT-like processes ^36^. Consistently, STAT3 inhibition was shown to reduce invasion and migration of cells ^12,37^. Since Stat3^Δ^ cells display a MITF-driven phenotype, it is interesting to note that HDAC inhibitors (HDACi) like Panobinostat have been shown to suppress MITF ^38^. Suppression of MITF activity by HDACi was pioneered as therapy for melanoma patients with high levels of MITF expression ^39^. HDAC inhibitors Panobinostat and Vorinostat are currently being tested as adjuvant therapy (NCT02032810), or as stand-alone treatment (NCT02836548) after BRAF^V600E^ inhibitor resistance (data from clinicaltrials.gov).

CEBP family members are well established downstream targets of STAT3 ^40,41^. Additionally, CEBPa/b/d DNA binding sites were reported in the murine *Mitf* promoter ^42^. Loss of STAT3 diminished expression of CEBP family members and CEBP binding motifs were less accessible as determined by ATAC-seq at the *Mitf* locus. Importantly, expression of CEBPa/b was sufficient to repress MITF, similar as reported in myeloid cells ^42^. Our study highlights the prognostic significance of viewing high or low levels of MITF or STAT3 and CEBP transcription factors in the etiology and progression of melanomas. Interestingly, when *STAT3, CEBPA, CEBPB* and *MITF* are used as criteria to split patient data into a low and high risk group, the low risk group reassembled the *STAT3, CEBPA, CEBPB* high and *MITF* low group, while the high risk group fits to the expression data resembling the *MITF* high expressing cohort. These findings are supported by studies which identified an impact of the proliferative status of melanoma on patient survival by quantifying Ki67, Proliferating Cell Nuclear Antigen (PCNA) and Cyclin D1 ^43,44^. This indicates, that patient survival times in melanoma are not determined by the degree of invasiveness, but rather by the amount of proliferation.

Our findings imply that STAT3 is a vulnerable node for fighting melanoma migration and invasion. We provide a rationale for clinical exploration of MITF inhibitors in combination with STAT3 inhibition, since MITF high expression is associated with poor survival. Importantly, here we identified STAT3 as a critical regulator of the MITF pathway, which has been recently shown to be the key pathway activated after acquired resistance to mitogen-activated protein kinase (MAPK)-targeted therapy ^45,46^. We conclude that a balanced YIN/YANG expression of STAT3/MITF controls melanoma fate and we propose that the main role of STAT3 is to promote invasion and metastatic spread.

## Materials and Methods

### Animals and generation of transgenic mice

Mice carrying the *Stat3*^*floxed*^ allele ^22^ were crossed with transgenic mice carrying melanocyte-specific expression of the *NRAS*^*Q61K*^ oncogene in an INK4A-deficient background ^21,47^ and tyrsosinase *Cre* ^48^. Human *NRAS*^*Q61K*^ and *Cre* are targeted to the melanocyte linage by tyrosinase regulatory sequences. Compound *Tyr∷NRAS*^*Q61K*^ *Ink4a*^*-/*-^ ; *Stat3*^*flox/flox*^*; Tyr∷Cre* mice were termed Stat3^Δ^ for simplified reading and maintained on a C57BL/6Jx129/Sv background. In all experiments described, littermates lacking *Tyr∷Cre* were used as controls termed Stat3^fl^. Immunocompromised NOD.CgPrkdc^scid^ Il2rg^tm1Wjl^/SzJ mice (NSG) recipients were purchased from Harlan. For all analyses male and female mice were used and analyzed in gender-specific ways.

### Xenograft experiments

A total of 1×10^6^ cells were injected into the left and right flank of 8–10 weeks old NSG mice. Tumor volumes were evaluated every 48 hours by measuring two perpendicular diameters with calipers. Tumor volume was calculated using the following equation: (width*width*length)/2. The experiment was stopped when tumors exceeded a calculated size of 600 mm^3^.

### Cell lines

Mouse melanoma cell lines were established from lymph nodes of diseased mice (25-30 weeks of age) by single cell dissociation and culturing. Multiple individual primary pools of cell lines per Stat3^fl^ and Stat3^Δ^ melanoma cell genotype were generated, upon which we selected two pools each for further detailed analysis. Human *BRAF* mutated cell lines WM793B, 451Lu and WM35 were bought from ATCC, all three devoid of INK4A proteins. The human *NRAS* mutated cell line WC125 (WM3854) was bought from the Coriell Institute (Wistar Collection) also devoid of INK4A proteins. All cell lines were cultivated under standard conditions (95% humidity, 5% CO_2_, 37°C) and maintained in standard medium (DMEM) supplied with 10% Fetal Bovine Serum (FBS), 10 U/ml Penicillin, 10 µg/ml Streptomycin, 2 mM L-Glutamine (all from Gibco) and 2 µg/ml Ciprofloxacin (Sigma). Cell stimulations were performed with human OSM (20 ng/ml), human IL-6 (20 ng/ml) or murine IL-6 (20 ng/ml; all from Immunotools). Murine OSM (20 ng/ml) was purchased from R&D. For transfection of murine melanoma cells, the cells were seeded in 6-well plates and starved overnight in 2% DMEM without antibiotics. Cells were transfected with 2 µg CEBPa, CEBPb or STAT3ER^T2^ plasmid using lipofectamine 3000 (Invitrogen) according to the manufacturer’s protocol and analyzed after 24 hours. After transfection, the cells carrying the STAT3ER^T2^ plasmid ^24^ were selected for four weeks in 1 µg/ml puromycin (Sigma) and expression was tested by Western blot (Fig. S3 C).

### 3D proliferation and *in vitro* migration assays

2500 melanoma cells were seeded in a 96 well round bottom plate and the diameter of formed spheroids was evaluated with the ImageJ software after 72 hours (Rasband, W.S., ImageJ, US National Institutes of Health, Bethesda, Maryland, USA, http://imagej.nih.gov/ij/). 30 spheroids, each consisting of 2.500 melanoma cells were formed in a 96 well round bottom plate. After four days of formation the spheroids were embedded into a 3D rattail-collagen matrix and grown in DMEM for 20 hours. Invasion was evaluated as described by ^49^. For the *in vitro* cell migration assay 3×10^4^ cells were seeded in serum free DMEM onto 8 µm pore transwell inserts (BD Biosciences). As attractant DMEM containing 10% FBS was used. Cells were allowed to migrate 8 hours, then the cells were fixed with 4% formaldehyde, the cells on the upper side of the transwell were wiped away and migrated cells were stained with 0.1% crystal violet (all from Sigma). Six fields of view were taken with microscope images using a 10x magnification with cell counting.

### RNA interference and lentiviral transduction

RNA interference and lentiviral transduction experiments were performed as described in ^50^. The following shRNA constructs selected from the Mission TRC shRNA library (Sigma) were used: shRNA STAT3 (TRCN0000071456) and scrambled control shRNA (SHC002). Transduced cells were selected for puromycin resistance prior to further analysis. The functionality of shRNAs was validated by realtime-PCR and Western blot analysis using antibodies listed in Table S3.

### RNA isolation and real-time PCR

RNA was isolated by TriZol (Life Technologies) extraction. Complementary DNA transcription was performed by RevertAid™ H Minus First Strand cDNA Synthesis Kit (Fermentas) according to the manufacturer’s instruction. Primer pairs are listed in Table S1. Real-time PCR was performed in triplicates on a RealPlex Master-cycler (Eppendorf). Values were normalized to either murine or human ACTB mRNA expression. Results were quantified using the Delta C(T) method ^51^.

### Western blotting

Protein lysates, SDS–PAGE and Western blotting were performed according to standard protocols. Nitrocellulose membranes (Amersham) were incubated with antibodies listed in Table S3. Semi-quantitative densitometric analysis of bands was performed with the ImageJ software according to standard procedures.

### Melanin absorption assay

Absorption of Melanin in mouse melanoma cells was measured at 410 nm in an Infinite Nanoquant photometer (Tecan) using the supernatant of 5×10^5^ cells 48 hours after seeding. Two biological replicates with six independent measurements were performed.

### Immunohistochemistry, immunofluorescence and tissue microarrays

Mouse tissue was fixed overnight in 4% phosphate buffered formaldehyde solution, dehydrated, paraffin embedded and cut. 3-µm-FFPE tissue sections were stained with Hematoxylin (Merck) and Eosin G (Roth). Immunohistochemistry and immunofluorescence was performed using antibodies listed in Table S2. For immunohistochemical analysis of cutaneous mouse melanomas, tissue microarrays were built including 28 cutaneous melanomas and 28 positive lymph nodes, each represented by duplicate core biopsies. Normal skin and lymph nodes served as controls. For quantification, high-power field sections (x200 objective) were obtained for each sample and analyzed with HistoQuest (TissueGnostics) (Fig. 1 B and C, Fig. S1 B) quantification software or Tissue Studio Software^®^ (Definiens) (Fig. 4 B and 7 C) as described ^52,53^. In brief, cut-offs (to differentiate between positive and negative cells) were set for all samples and the resulting positive intensities were normalized to Hemalaun staining intensity. Images were taken with a Zeiss Imager Z1 microscope. Immunofluorescence images were taken with a Leica TCS SP8 Microscope.

### Immunocytochemistry

To characterize murine cells as melanoma cells, they were trypsinized and 2×10^5^ cells were centrifuged using the Thermo Scientific Shandon Cytospin (Shandon Scientific). The samples were fixed in methanol (10’, -20°C) and dried for one hour at room temperature. Samples were stained by S100b by standard methods (Fig. S1 B, Table S2). For immunofluorescence staining on murine melanoma cells, 1.5x10^5^ cells were seeded on 4-well chamber slides (NUNC). Cells were grown in Lab-Tek^®^ 4-well chamber slides. Cells were fixed in 4% phosphate buffered formaldehyde solution and stained using antibodies from Table S2.

### Whole transcript expression profiling, and gene-set enrichment analysis

Total RNA was isolated using the RNAeasy Kit (Qiagen), pooled and hybridized to GeneChip Mouse Gene 1.0 ST array (Affymetrix). 24 hybridizations were performed (see GEO submission) comprising 12 replicates from each genetically defined cell line (two cell lines per Stat3^fl^ or Stat3^Δ^ murine melanoma cell line group including individual cytokine stimulation with either IL-6 or OSM). RNA was isolated from cells that were either unstimulated (full media without cytokine) or stimulated with fresh media containing either 5 ng/ml murine IL-6 or 10 ng/ml murine OSM for 2 hours. The command-line version of GSEA was used for gene-set enrichment analysis (GSEA) ^54,55^, with normalized log2 transformed expression values as input and “Signal2Noise” as ranking metric. Further non-default settings were -collapse false, - permute gene-set, and -set_min 3. In case of gene-sets with human gene ids, murine gene symbols were converted with the help of an orthologue list downloaded from the Ensembl gene browser web site (v75; GRCh37.p13). Heatmaps show hierarchical clustering of log-transformed gene expression values (Euclidean distance metric). Gene-sets were compiled from literature. Whole genome expression data was deposited at GEO (GSE95063).

### ATAC-seq

Chromatin accessibility mapping was performed using the ATAC-seq method as previously described ^56^ with minor adaptions. Duplicates of two different Stat3^fl^ and Stat3^Δ^ cell lines were used for the assay. A volume containing 50.000 cells were harvested with the aid of an automated cell counter (TC20 Bio-Rad) and spun at 500g for 5 minutes at 4°C in PBS. After centrifugation, the pellet was carefully resuspended in the transposase reaction mix (12.5 µl 2xTD buffer, 2 µl TDE1 (Illumina), and 10.25 µl nuclease-free water, 0.25 µl 1% Digitonin (Sigma)) for 30 min at 37 °C. Following DNA purification with the MiniElute kit eluting in 12 µl, 1 µl of the eluted DNA was used in a quantitative PCR reaction to estimate the optimum number of amplification cycles. Library amplification was followed by SPRI size selection to exclude fragments larger than 1,200 bp. DNA and concentration was measured with a Qubit fluorometer (Life Technologies). Library amplification was performed using custom Nextera primers ^57^. The libraries were sequenced by the Biomedical Sequencing Facility at CeMM using the Illumina HiSeq 3000/4000 platform and the 50-bp single-end configuration. Whole genome sequencing data was deposited at GEO (GSE119540).

### Preprocessing and analysis of ATAC-seq data

ATAC-seq reads were trimmed using Skewer ^58^ and aligned to the GRCh37/hg19 assembly of the human genome using Bowtie2 ^59^ with the ‘-very-sensitive’ parameter. Duplicate reads were removed using the sambamba ^60^ ‘markdup’ command, and reads with mapping quality >30 and alignment to the nuclear genome were kept. All downstream analyses were performed on these filtered reads. Genome browser tracks were created with the genomeCoverageBed command in BEDTools ^61^ and normalized such that each value represents the read count per base pair per million mapped and filtered reads. Finally, the UCSC Genome Browser’s bedGraphToBigWig tool was used to produce a bigWig file. Peak calling was performed with MACS2 ^62^ using the ‘-nomodel’ and ‘-extsize 147’ parameters, and peaks overlapping blacklisted features defined by the ENCODE project ^63^ were discarded. We created a consensus region set by merging the called peaks from all samples across cell lines, and we quantified the accessibility of each region in each sample by counting the number of reads from the filtered BAM file that overlapped each region. To normalize the chromatin accessibility signal across samples, we first performed quantile normalization using the R implementation in the preprocessCore package (‘normalize.quantiles’ function).

### Statistics

All values are given as means±standard error of the mean (s.e.m.) or standard deviation (s.d.) as indicated. Data were analyzed by GraphPad Prism^®^ 5. *In vitro* data, Western blots, RT-PCRs and viability assays were repeated at least three times. Scatter plots were calculated from the publically available GSE19234 and the TCGA study on cutaneous melanoma ^64,65^. RNA expression data sets were generated with SPSS v21. Pearson correlation analysis was performed to calculate the *P*-values of the graphs. Numbers of animals are stated in each figure or figure legend. Differences were assessed for statistical significance by an unpaired two-tailed t test and by the Log-rank (Mantel-Cox) test (for Kaplan-Meier plots). *P*<0.05 was accepted as statistically significant and P values are considered as follows: \**P*<0.05; \*\**P*<0.01; and \*\*\**P*<0.001.

### Study approval

Human tissue samples were collected after signed, informed consent was provided and approval for studies was obtained from Ethics Committee of the Medical University of Vienna, EK 405/2006, extension 11/10/2016. All mice were bred and maintained under standardized conditions at the Decentralized Biomedical Facility of the Medical University Vienna according to an ethical animal license protocol complying with the Austrian law and approved by the “Bundesministerium für Wissenschaft und Forschung” (BMWF-66009/0281-I/3b/2012).

## Acknowledgments

We thank Safia Zahma, Birgit Schütz, Eva Bauer, Michaela Schlederer and Karin Neumüller for excellent technical assistance. We thank Anna Ringler and Bernd Boidol for excellent assistance with the inhibitor screening library. We also thank the Biomedical Sequencing Facility at CeMM for assistance with next generation sequencing. VP was supported by the grant AIRC IG16930. RM and AS were supported by two network grants SFB-F061 and SFB-F047 and a private melanoma research donation from Liechtenstein. RS, KK, MV and MM were supported by P 25336-B13, all from the Austrian Science Fund (FWF).

## Author’s contribution

Authorship note: R. Moriggl and M. Mikula contributed equally to this work.

Initial design and work was performed by RM, MM and AS. RS, AS, MM and RM wrote the original draft of the manuscript. RS and AS designed, performed and analyzed experiments. MV performed *in vitro* 3D proliferation and invasion experiments. KK performed *in vitro* migration experiments and quantified xenograft and evaluated patient TMA stainings. BW, HTTP, RZ and BK helped designing experiments and provided major technical support. JH performed and analyzed xenograft experiments. CS and FA performed shRNA knockdown. DSc, AFR and TK analyzed *in silico* ATAC-seq or melanoma patient data. PP provided and stained human melanoma TMAs. SK and TK significantly assisted in data analysis. DSt, LK, LL, VP, FB, TY, SK, FA and MH reviewed and edited the manuscript. All authors have given approval to the final version of the manuscript.

